# Decoding antibody-antigen recognition: the BL8 language and its application in calibrating deep learning predictions

**DOI:** 10.64898/2026.10.30.763264

**Authors:** Tingwei Miao, Ruohan Zhu, Ruiqi Lin, Juncheng Qian, Yi Feng, Shuang Chen, Yuxin Wang, Daoyou Chen, Yanling Wu, Tianlei Ying

## Abstract

Antibodies are central to immunology and therapeutics, and understanding how they recognize antigens is essential to rational drug design. This recognition depends on the interplay of interface geometry, physicochemical interactions, sequence context, and complementarity-determining region (CDR) conformation. However, traditional analyses often oversimplify this complexity, which becomes particularly problematic when evaluating the massive and frequently overconfident structural hypotheses generated by modern deep learning models. Here, we introduce BL8, an interpretable, eight-view structural language that comprehensively parameterizes interfaces across topology, shape, interaction, CDR conformation, flexibility, interfacial area, amino-acid-pair composition, and sequence background. Using 15,396 experimental interface records from the Structural Antibody Database (SAbDab), we built a fixed reference atlas containing 1,498 discrete BL8 signatures. The atlas shows that natural interfaces occupy a diverse, long-tailed structural space in which similar geometric scaffolds can be associated with different chemical and sequence features. Furthermore, we projected 28,501 predicted complexes involving programmed death-ligand 1 (PD-L1), the SARS-CoV-2 receptor-binding domain (RBD), and programmed cell death protein 1 (PD-1), generated using AlphaFold 3 (AF3), Protenix-v2, and OpenDDE, into the reference space. Acting as a robust underlying calibration layer, BL8 successfully decoupled physical viability from native confidence illusions, revealing cross-target hallucinations and filtering out 15.3% to 20.1% of highly deceptive false-positive predictions. Crucially, by benchmarking distinct generative behaviors across models, it enabled the extraction of highly reliable, multi-model consensus out-of-distribution binding paradigms. Together, BL8 bridges foundational biophysics with large-scale computational predictions, providing an essential benchmarking framework for discovering novel mechanisms in the era of artificial intelligence.

**Highlights:** BL8 encodes antibody–antigen interfaces through eight interpretable views.

An atlas of 15,396 interfaces defines 1,498 experimental BL8 signatures.

BL8 reveals long-tailed diversity in antibody–antigen recognition.

BL8 rejects 15.3%–20.1% of high-confidence false-positive predictions.

Cross-model consensus prioritizes candidate binding modes beyond the atlas.

## 1. Introduction

Antibodies are widely used in immunology, diagnostics, and biomedicine. Their function depends on the ability of the variable-region paratope to recognize an antigen epitope with appropriate specificity, affinity, and selectivity [1, 2]. Understanding this recognition process is therefore central to antibody engineering, affinity maturation, and structure-based molecular design [3]. An antibody–antigen interface cannot be described by a single contact site or energetic score [4, 5]. Rather, it is a spatially organized surface shaped by topology, shape complementarity, packing, hydrogen bonding, electrostatics, hydrophobic contacts, solvent exposure, conformational adaptation, and sequence context [6, 7].

Structural biology has established the physical principles used to describe protein–protein recognition. Early studies emphasized shape complementarity, buried surface area, and the energetic contributions of interfacial contacts [8]. In antibody structures, the complementarity-determining regions (CDRs) form the principal recognition surface, but their contributions vary and depend on the surrounding framework and antigen context [9]. The concept of canonical structures provided a framework for organizing recurring CDR conformations, which were later refined through quantitative clustering [8, 10]. These findings indicate that CDRs are sufficiently constrained for structural comparison, but CDR identity alone does not describe the complete binding interface [11].

As structural datasets expanded, resources such as SAbDab and standardized numbering systems enabled systematic comparisons of paratope and epitope features across thousands of structures [12-14]. Large-scale contact analyses have quantified residue composition and hotspot patterns, while other studies have proposed compact interaction vocabularies and quantitative measures of complementarity [4, 5]. Together, these studies show that antibody–antigen recognition exhibits recurring structural preferences, but that these preferences overlap and depend on context rather than defining a single universal interaction pattern [1, 15, 16].

Despite these advances, the structural space of antibody–antigen interfaces remains difficult to characterize comprehensively. Experimental structures sample only a small and uneven portion of the diversity of antibody sequences, targets, and architectures. In addition, many analyses focus on one descriptive dimension, such as CDR conformation, contact composition, or shape complementarity [17]. These dimensions are related but not interchangeable: two complexes may have similar buried surface areas yet differ in contact topology, interaction chemistry, or flexibility. A useful representation should therefore integrate multiple views, remain traceable to measurable features, and avoid reducing recognition to a single affinity-related rule.

The urgency for such a multiview representation has grown with the rapid development of complex-structure prediction methods, including AlphaFold 3 and related deep-learning models [18, 19]. These methods can generate large ensembles of predicted antibody–antigen conformations, including for targets without experimental structures. However, generative models can produce physically implausible interfaces with high confidence scores. Native confidence metrics, such as predicted interface error, help assess a model’s internal consistency but do not independently determine whether an interface matches experimentally observed physical patterns. Conversely, a predicted interface absent from the experimental catalog is not necessarily incorrect: it may reflect undersampling, a rare target-specific interaction, or a novel binding mode outside the observed distribution (out-of-distribution, OOD) [20-22]. Evaluating large prediction sets therefore requires a fixed, multidimensional experimental reference space in which observed (covered) and unobserved (out-of-vocabulary, OOV) combinations can be assessed separately from native confidence scores.

Here, we introduce BL8, an interpretable eight-view structural language for antibody–antigen interfaces. From 15,396 interface records in SAbDab, we constructed a fixed reference atlas covering topology, shape and geometry, interactions, CDR conformation, flexibility and disorder, solvent-accessible interfacial area, amino-acid-pair contact composition, and sequence composition. The atlas contains 1,498 naturally observed BL8 signatures and provides a discrete description of structural diversity. We projected cross-target predictions for PD-L1, the SARS-CoV-2 RBD, and PD-1 from AlphaFold 3, Protenix-v2, and OpenDDE into this fixed reference space. This analysis allowed us to examine cross-target misattribution and compare the models’ distinct patterns of covered and OOV signatures. By treating OOV status as a structural descriptor and comparing predictions across models, BL8 also provides a route to identifying candidate out-of-distribution (OOD) binding modes supported by model consensus.

## 2. Results and discussions

### 2.1. The BL8 reference atlas defines a multidimensional structural coordinate system

To systematically decode the structural diversity of antibody-antigen recognition, we constructed a fixed and multidimensional BL8 reference atlas based on the SAbDab database [12]. We began with 10,779 Protein Data Bank (PDB) structures from the IMGT (International ImMunoGeneTics information system)-renumbered branch of SAbDab. During curation, we removed unrelated chains, non-protein ligands, ions, and bulk solvent while retaining crystallographic waters within 4.5 Å of both the binder and antigen to capture water-mediated interactions. This curation yielded 15,396 high-quality antibody–antigen interface records from 8,273 independent PDB structures (Fig. 1A), which, to our knowledge, constitute the largest analytical dataset of experimentally resolved antibody–antigen interfaces to date. Precise chain-role annotation allowed a single crystal structure to contribute multiple distinct interface records, preserving the observed diversity of conformations and binding modes. We defined candidate contacts using a 4.5 Å heavy-atom distance threshold and strict close contacts using a 3.5 Å threshold. CDRs were delineated using the Antigen Receptor Numbering and Classification (ANARCI) tool with instance-aware mapping to IMGT [23].

**Figure 1.**
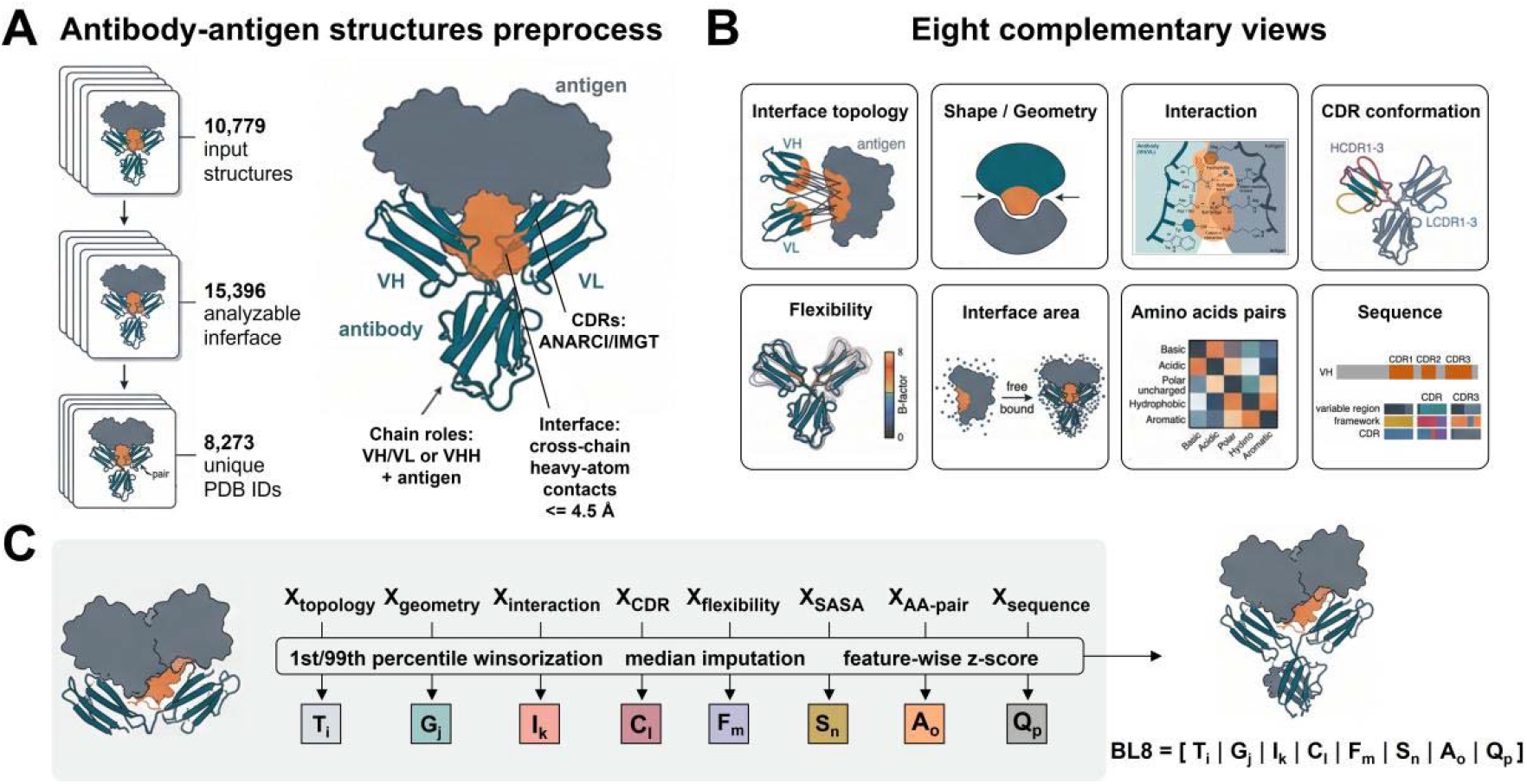
Construction pipeline of the multidimensional BL8 structural reference atlas. (A) Preprocessing and precise chain-role annotation of antibody-antigen structures from the SAbDab database. (B) Eight complementary views comprehensively characterizing the binding interface. (C) The workflow mapping 328 numerical features into discrete, complex-level BL8 state signatures.

Traditional studies are often limited to a single structural perspective, whereas BL8 introduces 328 numerical features to construct eight highly complementary interface dimensions (Table S7), enabling a panoramic characterization of the binding interface. This comprehensive feature set encompasses 230 foundational structural and interaction features, 64 amino acid property-pair features, and 34 sequence composition features. Specifically, interface topology delineates the spatial distribution of the contact network across the CDRs and framework regions; shape and geometry precisely quantify the contiguity, spatial packing, and surface complementarity of local patches, aligning with established interface research [6, 7]. Within the physicochemical interactions dimension, we systematically decoupled and quantified geometry-constrained hydrogen bond networks (encompassing direct and water-mediated hydrogen bonds calculated via explicit hydrogen addition) alongside distance-based salt bridges, hydrophobic, and aromatic contacts [24, 25]. Furthermore, by introducing principal component axis-based CDR conformation analysis, relative B-factor-based flexibility profiling, solvent-accessible surface area (SASA), amino-acid contact matrices matrices and variable-region sequence composition analyses 4, 5, 11, 17, 26], we achieved a deep integration spanning sequence, conformation, and interfacial physicochemical properties (Fig. 1B).

To translate this vast and complex feature space into a computable discrete representation, we subjected these eight dimensions to independent unsupervised clustering. Following rigorous 1st and 99th percentile winsorization, data imputation, and standard z-score scaling, KMeans clustering successfully extracted the core states within each dimension (Fig. 1C). By orthogonally combining the state codes from these eight dimensions, we ultimately and precisely identified 1,498 unique, naturally occurring BL8 signatures within the experimental data space. The resulting atlas provides an auditable coordinate system for antibody–antigen interfaces and a reference for evaluating large-scale structure predictions and candidate binding modes.

### 2.2. Multidimensional integration reveals a diverse, long-tailed, and complementary structural landscape

By utilizing this newly established multidimensional coordinate system, We first characterized the states defined within each view of the BL8 coordinate system (Table 1), and assessed their internal separation using silhouette coefficients (Fig. 2A). Flexibility/disorder view exhibited the highest score (0.543), indicating clear internal consistency, whereas topology (0.165), shape (0.155), and the amino-acid-pair matrix (0.077) displayed lower scores. Flexibility/disorder had the highest coefficient (0.543), whereas topology (0.165), shape (0.155), and the amino-acid-pair matrix (0.077) had lower values. These lower values are consistent with interfaces occupying a continuous structural manifold rather than forming fully separated biological classes [1, 4, 5, 17]. This pattern supports the concept of structural degeneracy [1, 15, 16]. Across 15,396 records, the six structural and physicochemical views (BL6) produced 185 observed state combinations (Fig. 2B). However, incorporating the amino acid property matrix and sequence background expanded this core repertoire to 1,498 unique BL8 signatures. This is visually corroborated in t-distributed stochastic neighbor embedding (t-SNE) projections, where topological states (Fig. 2C) demonstrate distinct spatial clustering, yet chemical states (Fig. 2D) overlap broadly within similar geometric regions. Complexes sharing identical contact areas or local geometries often achieve stable binding through entirely divergent chemical networks and sequence contexts [27, 28].

**Table 1.** The BL8 state dictionary defining distinct structural and physicochemical states across eight complementary interface views.

| BL8 view | State | n | Feature |
|---|---|---|---|
| Topology | HCDR3 concentrated | 3,529 | CDR/framework contact burden; active region |
|  | LCDR2 concentrated | 1,815 |  |
|  | Framework assisted | 2,249 |  |
|  | LCDR3 concentrated | 4,214 |  |
|  | HCDR1/HCDR2 platform | 3,589 |  |
| Geometry | Compact/contiguous | 6,626 | Patch radius, span, thickness, contiguity and segmentation |
|  | Broad/segmented | 8,770 |  |
| Interaction | H-bond sparse/mixed | 10,308 | H-bonds, salt bridges, hydrophobic/aromatic contacts and clashes |
|  | H-bond/electrostatic dense | 4,885 |  |
| CDR conformation | Extended structured LCDR3 | 3,133 | CDR contour, protrusion, curvature, planarity and secondary structure |
|  | Protruding curved LCDR1 | 1,888 |  |
|  | Compact or VHH-based | 10,375 |  |
| Flexibility | disorder | 1,277 | CDR-to-framework B-factor deviation |
|  | ordered | 14,119 |  |
| Interface area | Paired FV/exposed surface | 10,960 | SASA and buried surface area |
|  | Single-heavy/compact burial | 4,436 |  |
| Amino acids pairs | Aromatic-basic | 1,735 | Residue property contact fractions; CDR and all-interface |
|  | Polar-acidic | 2,814 |  |
|  | Opposite charge | 1,951 |  |
|  | Hydrophobic-aromatic | 2,211 |  |
|  | Basic-basic | 784 |  |
|  | Acidic-acidic | 403 |  |
|  | Polar-polar | 2,262 |  |
|  | Aromatic-hydrophobic | 1,768 |  |
| Sequence | Variable polar + CDR polar | 8,363 | IMGT variable/CDR/CDR3 composition and enrichment |
|  | HCDR3/CDR + variable | 5,570 |  |

**Figure 2.**
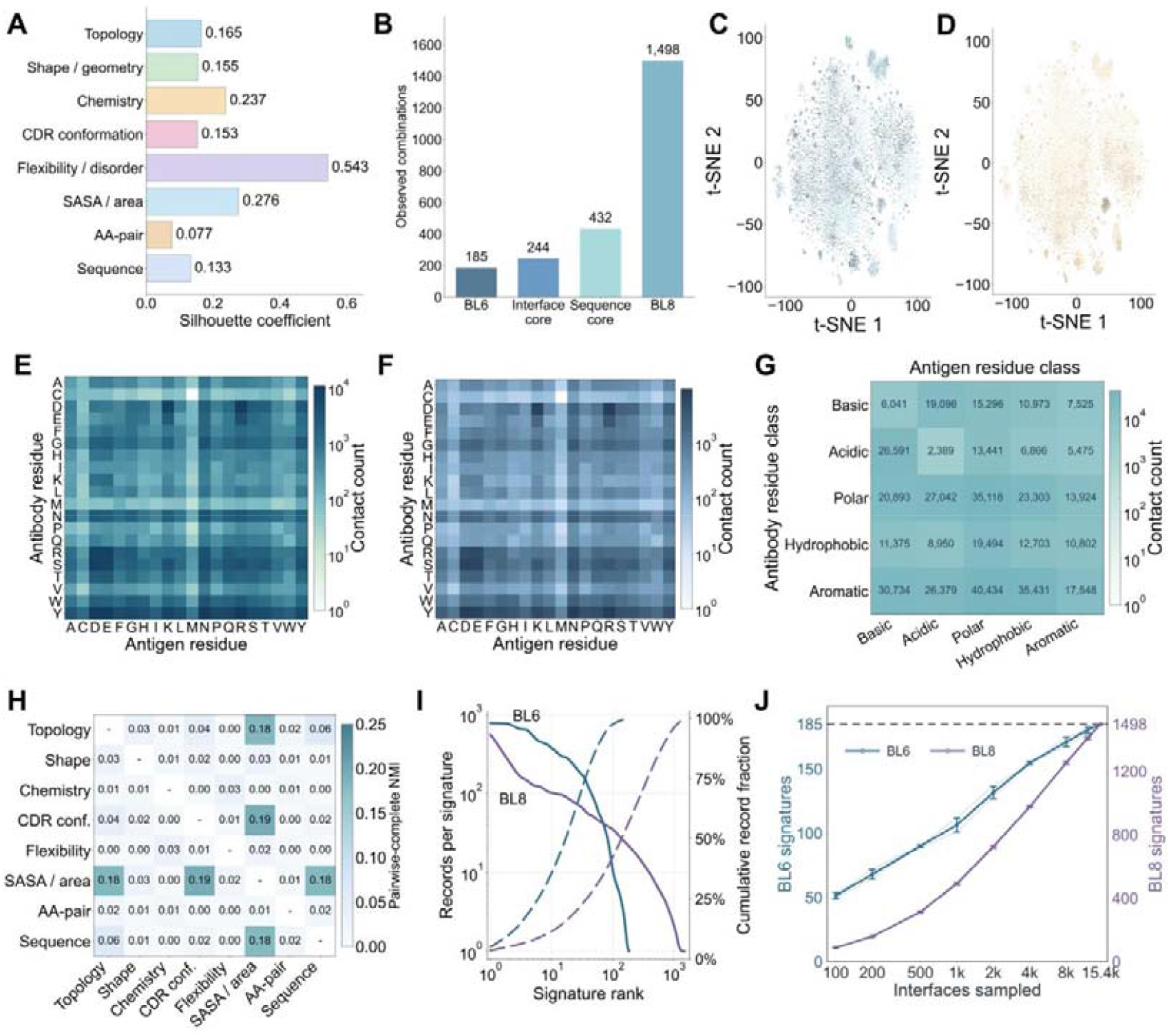
Multidimensional feature integration maps a highly diverse, long-tailed, and complementary structural landscape. (A) Silhouette coefficients evaluating the internal separability of each view. (B) Expansion of observed state combinations across increasing feature dimensions. (C, D) t-SNE projections of topology and interaction illustrating the continuous physical manifold of interface states. (E-G) Contact matrices demonstrating structural degeneracy and physicochemical versatility at the residue level. (H) Pairwise NMI revealing probabilistic couplings between views. (I) Rank-abundance distribution comparing purely structural (BL6) and fully integrated (BL8) signatures, demonstrating the pronounced long-tail characteristic driven by chemical and sequence diversity. (J) Random subsampling accumulation analysis evaluating the completeness of the current experimental structural database.

The contact matrices further decode this degeneracy at the residue level. In both the full-interface (Fig. 2E) and CDR-restricted (Fig. 2F) matrices, tyrosine (Tyr) participates in a broad range of frequent pairings, including Y–D, Y–R, Y–N, and Y–E, which remain common within CDRs. This pattern is consistent with the involvement of aromatic and polar residues in diverse paratope contacts [4, 15, 26]. At the property-class level (Fig. 2G), antibody-side aromatic residues formed enriched contact networks with antigen-side polar (40,434 occurrences) and hydrophobic (43,117 occurrences) residues. These metrics provide the physicochemical context that drives structural degeneracy, anchoring the BL8 signatures to concrete molecular interactions.

Despite this degeneracy, the eight views provide complementary rather than redundant information. We calculated normalized mutual information (NMI) for all 28 pairwise combinations (NMI) (Fig. 2H). The mean NMI was approximately 0.032, and 24 pairs had values below 0.05. The strongest associations were between CDR conformation and SASA (0.194), and between topology and SASA (0.182). These patterns are consistent with studies linking CDR conformation, the variable heavy (VH) and variable light (VL) environment, and interfacial burial [29-32]. The co-occurrence network (Fig. S6) illustrates these probabilistic relationships: some states co-occur more often than expected, but no single view determines the states of the others.

Having established the physical foundation of individual views and their probabilistic couplings, we next conducted a global analysis of the entire interface landscape to evaluate its overall distribution (Fig. 2I). Comparing rank-abundance curves for BL6 and BL8 showed that the long tail was present in both representations but was more pronounced in BL8. In BL6, the most frequent signature accounted for 5.09% of records, the ten most frequent signatures accounted for 35.70%, and 18 signatures covered half of the dataset. In BL8, the corresponding proportions were 3.59% and 11.49%, and 140 signatures were required to cover half of the records. Rare signatures, defined as those observed five times or fewer, increased from 54 in BL6 (29.19% of catalog signatures and 0.91% of records) to 888 in BL8 (59.28% of catalog signatures and 13.22% of records); the number of singletons rose from 19 to 363. Thus, the structural scaffolds represented by BL6 are more concentrated, whereas adding chemical and sequence features reveals a more combinatorially diverse distribution. From a biological perspective, antibody repertoire studies and structural consensus analyses both suggest that conserved structural scaffolds can coexist with highly diverse local chemical and sequence combinations [1, 15, 16, 26].

Beyond the static distribution, we also asked one of the most pressing questions in the field: has the current experimental database captured the complete universe of antibody-antigen binding paradigms, or will the continued addition of experimental data eventually converge to a finite set of structural languages? By conducting a random subsampling accumulation analysis (Fig. 2J), we observed that the BL6 structural catalog demonstrates a clear slowing trend, indicating that the purely geometric and topological macro-scaffolds are gradually approaching convergence. By contrast, the complete BL8 catalog exhibits a linear, non-converging trajectory as the sample size increases. These results indicate that the currently resolved structures are far from reaching the full biological repertoire of interface diversity. Rare signatures in the reference catalog therefore provide a useful baseline for evaluating target-specific diversity and the range of binding modes explored by structure-prediction models.

### 2.3. Physical interpretation of BL8 states

After establishing the diversity and distribution of the BL8 dictionary (Table 1), we examined the physical and biological properties represented by its states. To ensure that these mathematical clusters are not merely algorithmic artifacts, we mapped each state to concrete structural properties. The state inventory (Fig. 3A) clearly reveals the unevenness and diversity of interface architectures within the natural antibody repertoire. For example, interface topology spans five biological states, dominated by LCDR3 concentrated (n = 4,214) and HCDR3 concentrated (n = 3,529), confirming that no single paratope region universally serves as the sole anchor for antigen recognition. Furthermore, by conducting joint cross-analyses of these interpretable states, we elucidated the decoupled relationship between macroscopic interface geometry and local organizational modes. The joint distribution of topology and shape (Fig. 3A) demonstrates that an identical contact topology can map to entirely distinct geometric morphologies, which aligns with the understanding that CDR conformations and interface organizations exhibit combinatorial diversity in antibody structures [10, 11, 33]. Taking the HCDR3 concentrated topology as an example, it can form both highly compact, closed interfaces (n = 1,130) and broad, segmented contact networks spread across the antigen surface (n = 2,399). Quantitative analysis of these macroscopic states unveils a deeper physical principle: under the same topology, broad/segmented morphologies are typically accompanied by significantly larger interfacial burial areas (mean BSA). For instance, within the framework assisted topology, broad interfaces achieve a mean BSA of approximately 952 Å^2^, compared to just 651 Å^2^ for their compact counterparts. This directly proves that “which antibody regions bear the contact burden” and “how the contact surface unfolds in space” are two mutually independent physical dimensions. Relying solely on BSA or a single topological metric is fundamentally insufficient to accurately reconstruct the true architecture of an interface [34, 35].

**Figure 3.**
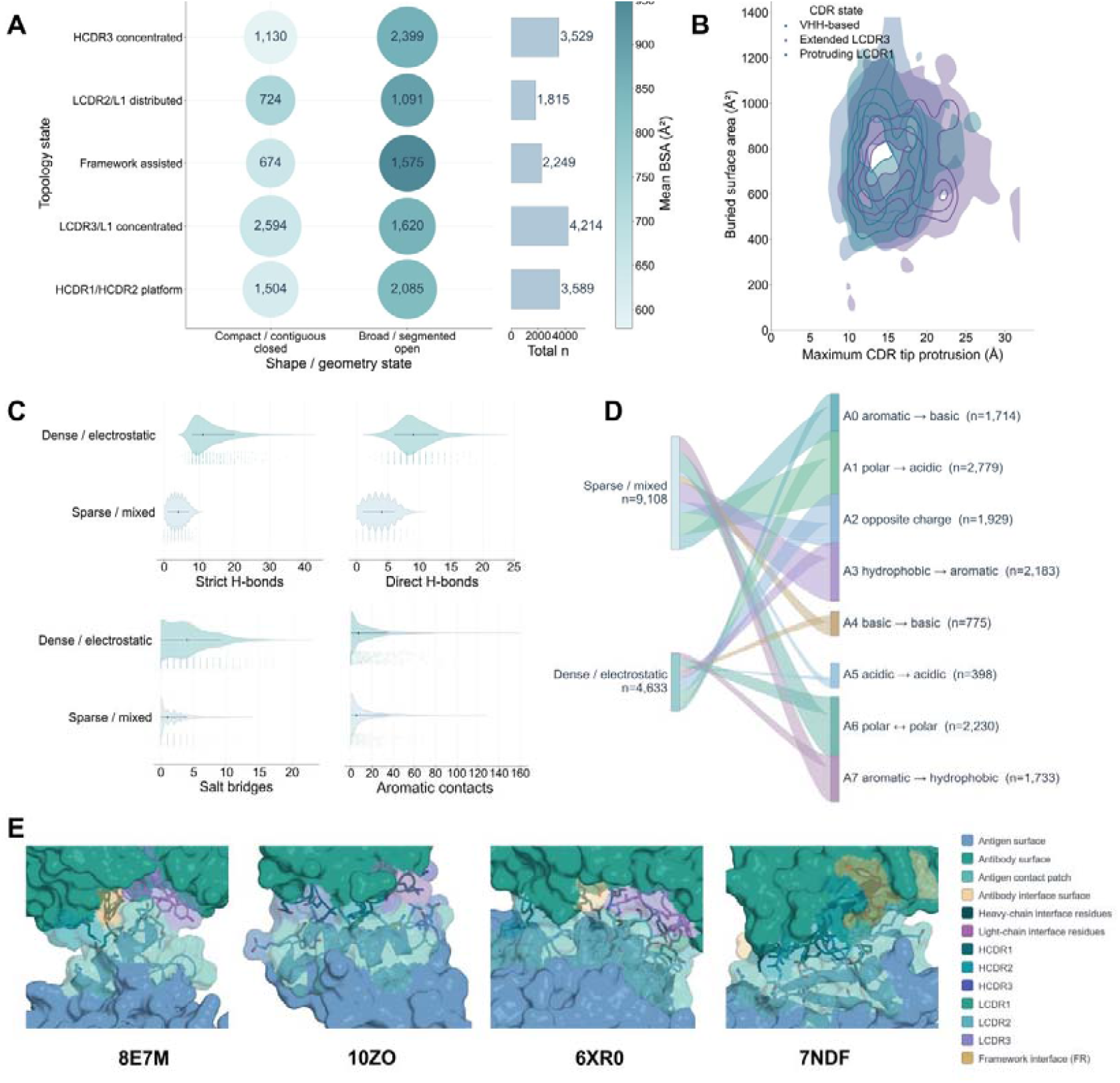
Structural interpretability and physical mapping of the BL8 state dictionary. (A) Joint distribution of interface topology and shape/geometry states decoupling macroscopic morphologies from local organizations, with marginal bar charts displaying the state distribution inventory. Bubble size indicates the number of observed interfaces (n), and color intensity represents the mean BSA. (B) 2D kernel density estimation of maximum CDR tip protrusion versus buried surface area (BSA). (C, D) Violin plots of physicochemical interactions and a Sankey diagram mapping macroscopic chemistry states to underlying amino-acid-pair compositions (right). (E) Representative 3D structural exemplars (8E7M, 10ZO, 6XR0, 7NDF) mapped to distinct BL8 binding paradigms.

We next examined CDR conformation in relation to interface burial (Fig. 3B). The two-dimensional kernel-density projection challenges the assumption that longer CDRs necessarily produce larger contact areas. CDR conformation depends on loop length, framework environment, and VH–VL pairing, and therefore cannot be described by length alone [10, 11, 29, 30, 33]. The three principal CDR-conformation states overlap across the continuous space defined by maximum CDR-tip protrusion and BSA. Most structures fall within a central region with protrusions of 10–20 Å and BSA values of 400–900 Å^2^; protrusions greater than 20 Å do not necessarily coincide with larger interface areas. This indicates that the actual binding scale is deeply constrained by CDR spatial orientation, multi-loop cooperativity, framework involvement, and the local geometric curvature of the antigen. The interaction and residue-property views also show why hydrogen-bond counts alone do not describe binding strength [6, 7, 24, 25]. In the violin plots (Fig. 3C), the H-bond/electrostatic-dense state (n = 4,885) had higher numbers of strict and direct hydrogen bonds and salt bridges. The H-bond-sparse/mixed state (n = 10,308), by contrast, showed greater reliance on hydrophobic and aromatic contacts across its long-tailed distribution. The Sankey diagram (Fig. 3D) shows a many-to-many mapping between these two broad interaction states and amino-acid-pair (AA-pair) compositions. The electrostatic-dense state was associated with several residue-pair patterns, including opposite-charge and polar–acidic pairings. The mixed state likewise mapped to multiple combinations, including hydrophobic–aromatic pairings. This multidimensional chemical representation precisely preserves the chemical redundancy and combinatorial diversity essential for antigen recognition.

To validate the structural fidelity of the complete BL8 state dictionary, we projected the state labels onto four representative three-dimensional complexes (Fig. 3E). The examples illustrate distinct binding modes: 8E7M has a compact, contiguous interface dominated by the light chain, with LCDR1 and LCDR3 forming a closed pocket; 10ZO has a single-domain antibody interface in which an HCDR3-concentrated topology protrudes into an antigen cleft; 6XR0 has a broad, platform-like interface involving HCDR1 and HCDR2 and a dense electrostatic network; and 7NDF has a framework-assisted interface in which framework residues (shown in gold) extend the contact surface alongside the CDRs. These exemplars provide compelling evidence that BL8 is not a set of mutually exclusive, rigid classifications, but rather a continuous coordinate system possessing both high-dimensional compression capability and exceptional physical interpretability, offering a unified analytical benchmark for subsequent evaluations of target-specific predictions or OOV interfaces beyond the known coverage scope [36, 37].

### 2.4. Cross-target evaluation of predictions for PD-L1, RBD and PD-1 using BL8

With the structural interpretability and physical fidelity of the BL8 reference space validated on experimental data, we applied this universal coordinate system to evaluate the massive structural hypotheses generated by current deep learning-based models. Considering the inherent inaccuracies and uncertainties of existing AI models in predicting antibody structures, we employed AF3, Protenix-v2, and OpenDDE to collaboratively generate predictions, thereby mitigating single-model biases as much as possible [18, 38]. We obtained target sequences from patents and directly projected 28,501 positive predicted structures of PDL1, RBD, and PD1 generated by the three models into this frozen reference space. This projection process did not involve any target-specific retraining or state redefinition, thereby ensuring the absolute objectivity and consistency of the evaluation criteria.

When evaluating these predicted complexes, relying solely on native model confidence metrics (such as ipTM, PAE, or GPDE) presents significant limitations. Recent benchmark studies have indicated that the score scales and interface error characteristics of different methods are not entirely comparable [19]. Density distribution curves reveal that the ipTM scores for AF3, Protenix-v2, and OpenDDE across different targets exhibit highly multimodal and target-dependent distribution patterns (Fig. 4A). Furthermore, even for high-confidence structures with similar ipTM scores, their cumulative distributions of interface errors in terms of PAE interaction or mean chain-pair GPDE differ drastically (Fig. 4B). More crucially, the evaluation consistency among models for the same input sequence is only moderate: the Pearson and Spearman correlation coefficients for ipTM and interface rank across the three models primarily fluctuate between 0.41 and 0.65 (Fig. 4C, Fig. S13). This moderate level of consistency profoundly indicates that different models likely rely on entirely distinct structural priors or sampling strategies. A structure deemed high-quality by one model might score extremely poorly in another. Therefore, native scores act as model-specific relative conditional probabilities and cannot be directly compared across models. This underscores the urgent need to introduce BL8 as an independent and interpretable structural language coordinate to objectively resolve which binding paradigm these high-scoring predictions physically belong to.

**Figure 4.**
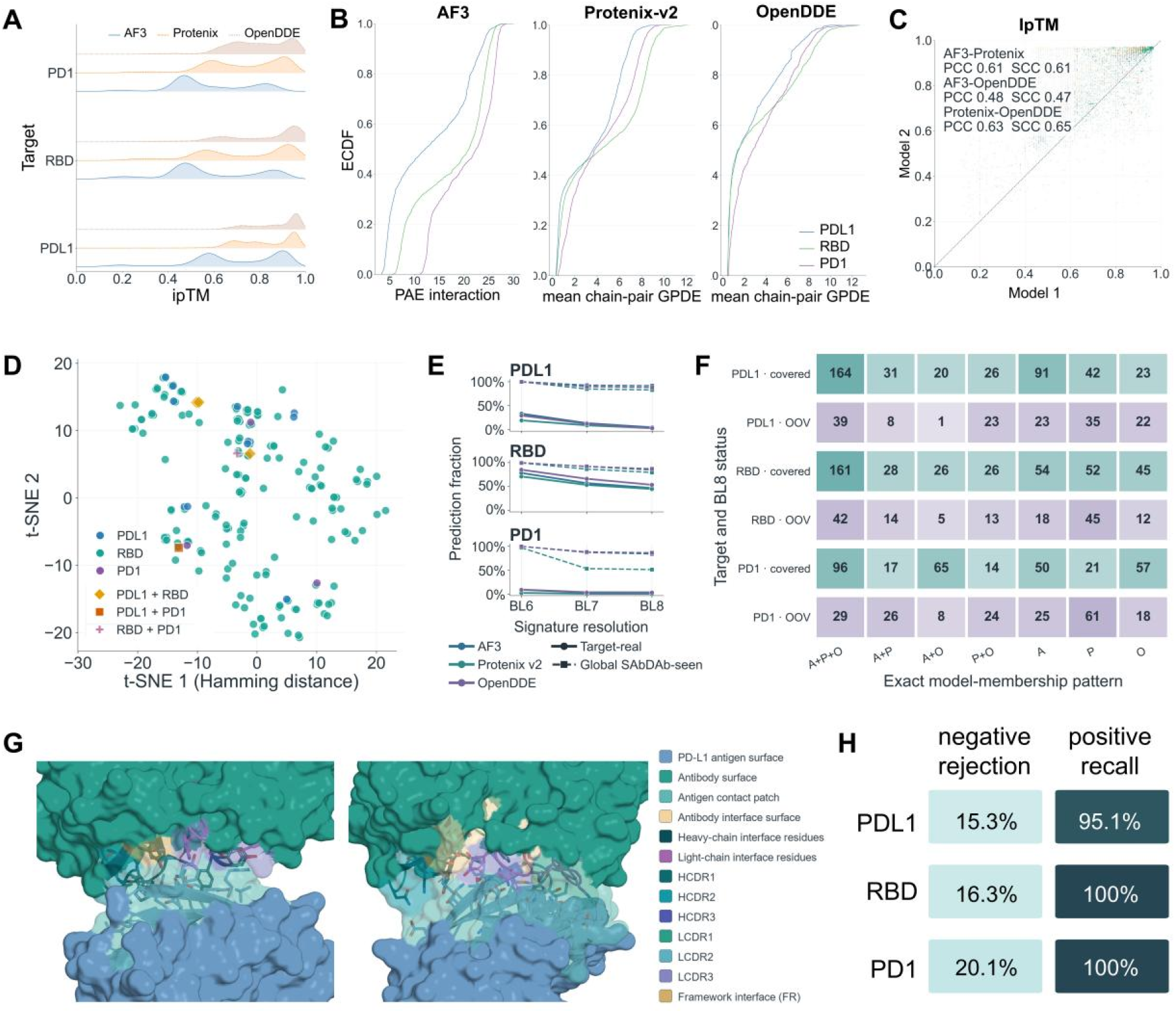
Cross-target evaluation and structural calibration of AI-predicted complexes using the BL8 structural language. (A) Density distributions of native confidence metrics (ipTM) across AF3, Protenix-v2, and OpenDDE for different targets. (B) Cumulative distributions (ECDF) of interface errors (PAE interaction or mean chain-pair GPDE). (C) Cross-model evaluation consistency for ipTM. (D) t-SNE projection mapping experimentally resolved SAbDab structures for PDL1, RBD, and PD1 into the BL8 manifold. (E) Prediction coverage evaluating the intersection with global SAbDab-seen and target-real structural spaces across varying resolutions (BL6 to BL8). (F) Exact model-membership patterns analyzing covered and OOV signature attribution across models. A: AF3, P: Protenix-v2, O: OpenDDE. (G) 3D structural representations of two of the most reliable multi-model consensus OOV binding paradigms for PDL1. (H) Statistical evaluation of BL8 acting as an underlying calibration layer for rejecting false positives (negative rejection) and recalling true predictions (positive recall).

To uncover the root cause of this prediction inconsistency, we first mapped the existing experimentally resolved structures for these three targets from the SAbDab database into the BL8 manifold (Fig. 4D). The experimental data distribution exhibits extreme target imbalance: there are 185 exclusive BL8 signatures for RBD, compared to a mere 12 for PDL1 and 3 for PD1. This severe scarcity of target-specific training data inevitably restricts the learning capacity of structural models. We subsequently analyzed the predicted structures by evaluating their intersection with both the global SAbDab-seen space and their respective target-real spaces across varying resolutions (Fig. 4E). A striking dichotomy emerged: all three models demonstrated exceptionally high coverage within the global structural space (particularly at the macroscopic BL6 resolution), indicating that they have successfully learned the generalized, global rules of antibody-antigen binding. However, their coverage of the target-specific space dropped precipitously for data-poor targets like PDL1 and PD1, maintaining acceptable levels only for the data-rich RBD (Fig. 4E). This discrepancy exposes a fundamental flaw in purely data-driven architectures: while they effectively interpolate global structural distributions, they systematically fail to capture the highly specific paratope-epitope recognition paradigms required for novel or data-scarce targets. Considering the distinct binding modes and specific paratope-epitope interaction preferences inherent to different targets, relying on a global understanding risks misattributing a binding mode from one target to another. Such cross-target hallucinations can lead to fatal failure rates in actual wet-lab validations, even though the generative models assign these predictions deceptively high native confidence scores.

Beyond assessing the coverage of known experimental states, the BL8 manifold allows us to investigate predicted signatures falling entirely outside the established global reference space, defined as OOV states. Signature attribution pattern analysis (Fig. 4F) reveals distinct generative behaviors and diversity tendencies among the models. AlphaFold 3 exhibits a highly conservative tendency, generating a large number of model-exclusive covered states (e.g., 91 for PDL1, 54 for RBD) while maintaining relatively low exclusive OOV states, reflecting its strong inductive bias toward memorizing known PDB templates. Conversely, Protenix-v2 demonstrates a highly exploratory behavior, producing the highest number of single-model exclusive OOV signatures across all targets (35 for PDL1, 45 for RBD, 61 for PD1). This stark contrast in covered/OOV ratios establishes BL8 not only as a structural filter but also as a powerful benchmarking suite to deeply evaluate the true generative capabilities, conservative biases, and exploratory boundaries of different AI models. While the majority of isolated single-model OOV states likely originate from prediction errors or local conformational artifacts, multi-model consensus provides a rigorous filtering mechanism. For instance, the three models collectively produced 39 unanimously agreed-upon OOV signatures (A+P+O) for PDL1, 42 for RBD, and 29 for PD1. Fig. 4G specifically visualizes two of the most reliable PDL1 interface paradigms extracted from this consensus OOV data. Because these structures were commonly predicted to have the same OOV signature by three architecturally independent models, this compellingly demonstrates that they represent a highly reliable, physically viable novel binding paradigm rather than mere model prediction noise. This in silico strategy, which unites the BL8 structural language with multi-prediction-model consensus, provides an extremely effective pathway for exploring and establishing reliable OOD binding modes that transcend existing wet lab crystallographic data. Given the current scarcity of high-resolution experimental structures and the inherent uncertainties and limitations of single deep learning model predictions, this approach can mine credible, entirely new binding architectures from massive prediction hypotheses, thereby offering invaluable data augmentation and mechanistic insights for discovery and engineering design in the antibody field.

More importantly, BL8 demonstrated robust capabilities as an underlying calibration layer for the aforementioned cross-target hallucination. We constructed the same amount of negative samples by combining cross-target antibodies-antigens. Fig. 4H clearly shows that BL8 maintains an exceptionally high positive structure retention rate (positive recall reaching 95.1%–100.0%) across all targets, meaning that stringent reference space constraints did not result in the loss of high-quality true predictions. When evaluating the negative rejection rate, the so-called negative structures we tested were actually false-positive structures judged as high-quality by the native metrics of these three structure prediction models. In other words, the native high confidence scores provided by these models were theoretically already erroneous. It is precisely under this stringent context of native model scoring failure that BL8, by examining the physical compliance of the structural language, still successfully corrected 15.3% to 20.1% of the models’ high-confidence prediction errors, effectively filtering out these highly deceptive pseudo-positive structures. In conclusion, BL8 provides a coordinate system that perfectly complements native model scores; it not only forcefully corrects the high-confidence biases of deep learning models but also accurately screens for reliable OOV candidate modes supported by multi-model consensus, opening a completely new evaluation dimension for the design and discovery of targeted molecules.

## 3. Conclusion

In this study, we developed BL8, an interpretable, multidimensional structural language for antibody-antigen interfaces. By systematically integrating 328 physical and sequence features—derived from the largest analytical dataset of experimentally resolved interfaces to date (15,396 records)—into eight orthogonal views, BL8 effectively transforms the highly degenerate and continuous interface manifold into 1,498 discrete, physically grounded signatures. This comprehensive reference atlas reveals that antibody-antigen recognition does not follow a single universal pattern but exhibits a pronounced long-tailed distribution. We demonstrated the universal prevalence of “structural degeneracy,” wherein conserved macroscopic geometric scaffolds synergistically couple with extreme combinatorial diversity in local chemical networks and sequence spaces.

Furthermore, BL8 exhibits exceptional physical interpretability, correcting the over-reliance on one-dimensional heuristics prevalent in traditional antibody design. Through joint cross-analyses, we elucidated the decoupled relationship between macroscopic geometric morphologies of interfaces and their local contact topologies.

Concurrently, this multidimensional perspective mitigates the bias of simplified structural proxies—such as assuming a simple linear correlation between CDR length and interfacial burial, or over-weighting raw hydrogen bond counts as the primary determinant of binding affinity. BL8 demonstrates that the actual binding scale and chemical complementarity are deeply constrained by multi-loop cooperativity, framework involvement, and high-dimensional physical redundancy, thereby providing a unified coordinate system to describe radically different binding paradigms.

Crucially, in an era where generative AI routinely produces massive and frequently “overconfident” structural hypotheses, BL8 provides an objective physical yardstick independent of native model metrics. When applied to evaluate cross-target predictions (PDL1, RBD, and PD1) from AlphaFold 3, Protenix-v2, and OpenDDE, BL8 successfully decoupled the true physical viability of interfaces from the confidence illusions generated by deep learning models. By exposing how purely data-driven architectures often force global binding rules onto specific, data-scarce targets, BL8 directly mitigates the risk of “cross-target hallucinations” that could otherwise lead to fatal wet-lab validation failures. Operating as a robust underlying calibration layer, it forcefully corrected high-confidence biases, effectively filtering out 15.3% to 20.1% of highly deceptive false-positive structures. More importantly, by benchmarking distinct generative behaviors—from highly conservative PDB memorization to aggressive structural exploration—it precisely extracted OOV states supported by multi-model consensus, enabling the in silico discovery of novel, OOD binding paradigms that transcend existing real-world crystallographic data.

In conclusion, as a highly auditable and robust structural benchmark, BL8 bridges the critical gap between foundational structural biophysics and large-scale computational hypotheses. It not only provides a solid theoretical foundation for systematically decoding the complexities of antigen recognition but also establishes an invaluable analytical framework for the artificial intelligence era. Moving forward, BL8 can serve as a standardized benchmarking suite to evaluate the true generative capacities and exploratory boundaries of emerging structural models, guide the rational design of target-specific antibody libraries, and accelerate the discovery of unprecedented therapeutic targeting mechanisms.

## 4. Materials and methods

### 4.1. Structure Preparation and Data Analysis

We constructed the reference atlas from experimentally determined antibody–antigen complexes in the SAbDab. To standardize residue numbering, we used the IMGT (International ImMunoGeneTics information system)-renumbered branch of SAbDab. The initial set comprised 10,779 Protein Data Bank (PDB) files. During curation, we removed unrelated chains, non-protein ligands, ions, and bulk solvent. We retained crystallographic waters located within 4.5 Å of both the binder and antigen. For residues with alternate locations, we kept the highest-occupancy conformer or, when occupancies were equal, the blank conformer. The unit of analysis was an explicitly annotated antibody–antigen interface record rather than the parent PDB file. Thus, a single asymmetric unit or multichain complex could contribute multiple records with distinct chain-role assignments. We required each interface to contain at least one valid local anchor patch, defined by at least ≥20 atom-pair contacts and a steric clash fraction of ≤0.05. The final frozen feature table comprised 15,396 records from 8,273 unique PDB identifiers. We delineated CDRs using instance-aware ANARCI-to-IMGT mapping.

### 4.2. Contact definition and geometric topology extraction

We defined residue contacts from cross-chain atomic interactions. For antibody atom a and antigen atom b, the candidate contact set P comprised pairs separated by no more than 4.5 Å between heavy atoms:

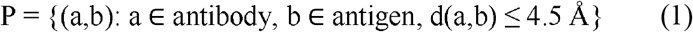

We defined strict close contacts, used to calculate atomic densities and construct the amino-acid-pair matrix, using a binary indicator and a 3.5 Å distance threshold:

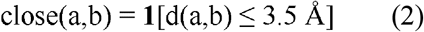

For each residue-pair row r, we aggregated the atomic contact burden, number of close pairs, and minimum and mean interaction distances:

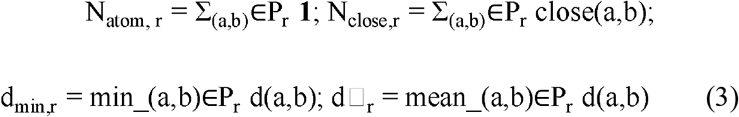

To characterize local geometry, we selected up to three antibody contact anchors per interface. The anchor score rewarded formal interactions and penalized steric clashes:

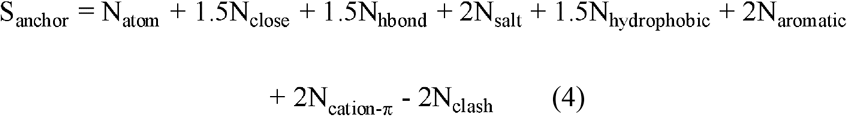

For each anchor, we defined a local patch with a 7.0 Å radius. We calculated patch morphology, including the radius of gyration (R_G_) and span, from the Cα coordinates of contacted residues:

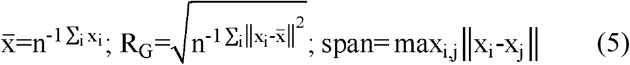

We derived shape descriptors, including linearity, planarity, and thickness, from the covariance eigenvalues (λ_1_≤λ_2_≤λ_3_) of the patch coordinates:

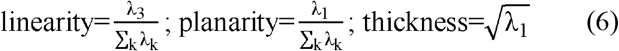

We quantified macroscopic structural descriptors, including surface closure, shape complementarity, packing fraction, and directional lock fraction. To describe interface topology, we distributed atomic contact burden across seven predefined regions (HCDR1–3, LCDR1–3, and framework) and calculated the topological entropy (H_topology_) and number of active regions (N_active_):

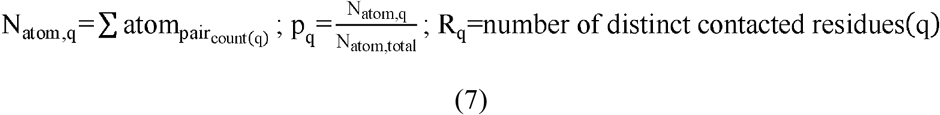

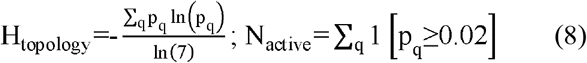

### 4.3. Interfacial chemistry and hydrogen-bond networks

We treated geometry-constrained hydrogen-bond networks separately from nondirectional physicochemical contacts. After adding explicit hydrogens with PyMOL’s h_add function, we evaluated candidate hydrogen bonds using relaxed and strict geometric criteria. A strict hydrogen bond, whether a direct protein–protein contact or a protein–water-mediated contact, was defined using donor (D), hydrogen (H), and acceptor (A) geometry:

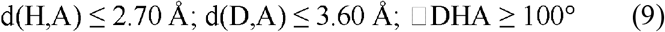

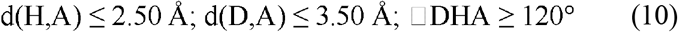

We also classified overlapping distance-based interactions among candidate atom pairs within 4.5 Å. The classes included salt bridges (≤4.0 between specified charged groups), hydrophobic contacts involving non-backbone carbon or sulfur atoms, aromatic contacts, cation–π interactions, and steric clashes (≤2.0). We normalized the relative density of each interaction class by the total close-contact burden:

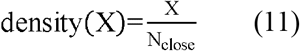

### 4.4. CDR conformation, flexibility, and solvent-burial metrics

We described the three-dimensional conformations of CDR loops using sequence-ordered Cα coordinates. Conformational descriptors included contour length (L), end-to-end distance (E), extension ratio, and a curvature proxy:

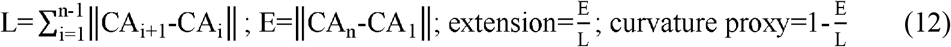

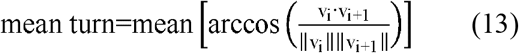

We derived loop shape descriptors (linearity, planarity, and thickness) using a principal-component-analysis (PCA)-like axis decomposition. Tip protrusion was defined as the maximum excursion from the axis connecting the first and last residues of each loop:

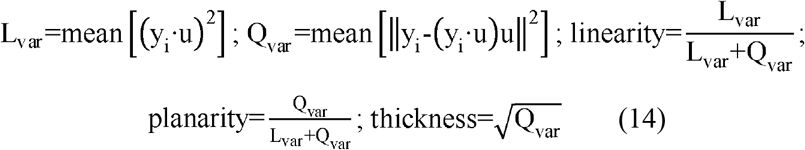

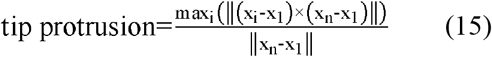

We estimated structural flexibility and local disorder from experimental Cα B-factors. For each CDR, we expressed relative mobility as a z score relative to the mean and population standard deviation (SD_B_) of the stable framework region:

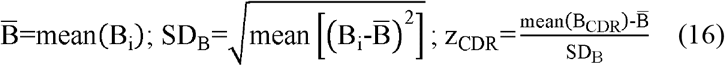

We calculated interfacial solvent burial using PyMOL’s dot_solvent algorithm. Solvent-accessible surface area (SASA) values for the free and bound states were used to derive absolute buried surface area (BSA) and the normalized fraction of antibody burial:

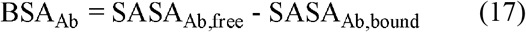

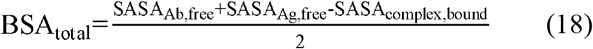

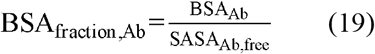

### 4.5. Amino-acid-pair matrix and sequence-composition encoding

To represent physicochemical compatibility while limiting sparsity among individual residue identities, we grouped standard amino acids into five exhaustive property classes: basic, acidic, polar uncharged, hydrophobic, and aromatic. For both the full interface and the CDR-restricted region, we calculated a 5×5 fractional matrix of residue-property contact pairs and the corresponding Shannon entropies:

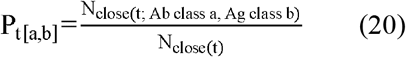

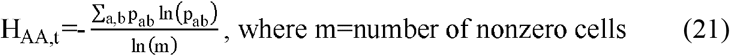

We also encoded sequence composition from the IMGT-mapped antibody variable domain. We calculated the enrichment of each residue class in the CDRs relative to the framework and the relative lengths of the loops to place the structural features in their antibody sequence context:

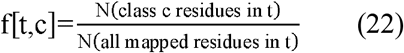

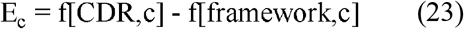

### 4.6. Independent view clustering and BL8 signature generation

The final interface representation comprised 328 features: 230 structural and interaction features, 64 amino-acid-pair features, and 34 sequence features. To limit the influence of extreme values, we winsorized continuous variables at the 1st and 99th percentiles:

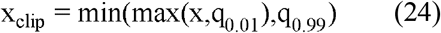

After median imputation, we standardized features using z scores calculated separately for each view:

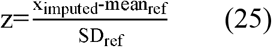

We clustered the eight views independently using K-means. For each view, we selected the number of clusters (K) primarily by maximizing the silhouette coefficient, using the Davies–Bouldin index as a secondary criterion. We concatenated the resulting state codes in a fixed order to form each complex-level signature:

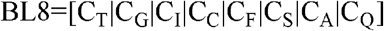

We assigned catalog identifiers only to complete state tuples observed among the 15,396 SAbDab reference records. This yielded a frozen reference space comprising 1,498 empirical binding paradigms.

### 4.7. AI structure prediction and candidate generation

To assess transferability of the BL8 framework, we generated positive antibody–antigen predictions for PD-L1, RBD, and PD-1 using three independent structure-prediction systems: AlphaFold 3 (AF3), Protenix-v2, and OpenDDE (Abag weight). For each model–record pair, we ran three independent random seeds, each producing five structural samples, for a nominal total of 15 structures. Before assigning BL8 states, we parsed each candidate’s model-native ranking score (e.g., ipTM) and all available confidence and interface metrics, including PAE interaction for AF3 and mean chain-pair global pairwise distance error (GPDE) for Protenix-v2 and OpenDDE. We selected the representative structure for each model–record pair using the model’s native ranking convention. BL8 therefore did not influence candidate selection and served only as an independent structural-language interpretation layer. In total, 28,501 positive representative structures were retained for cross-target analysis.

### 4.8. Fixed-space projection and model-membership assignment

We processed selected prediction structures with a harmonized prediction-side feature extractor. For observables unavailable in predicted structures, the extractor used proxies, such as heavy-atom donor–acceptor distance criteria in place of explicit water-mediated hydrogen bonds. We transformed the extracted features using the frozen SAbDab preprocessing parameters and assigned each structure to its nearest saved reference centroid by Euclidean distance; no target-specific reclustering was performed. We then compared the predicted state tuples with the frozen SAbDab catalog. Signatures matching one of the 1,498 empirical combinations were classified as covered, and unobserved combinations were classified as OOV. At the record level, we quantified model-to-model state transitions by tracking whether the same records were assigned covered or OOV states by different prediction systems. At the signature level, we assigned each unique predicted tuple a model-membership pattern (e.g., A+P+O for signatures observed in AF3, Protenix-v2, and OpenDDE). For quality-stratified analyses, we combined structural-language status with prespecified model-native confidence filters to assess structural patterns separately from internal confidence estimates.

## Code Availability

The BL6 and BL8 are available at https://github.com/SII-MTW/BL6-BL8-mapper.

## CRediT authorship contribution statement

**Tingwei Miao:** Writing – review & editing, Writing – original draft, Conceptualization, Investigation, Methodology, Formal analysis, Validation. **Ruohan Zhu:** Conceptualization, Investigation. **Ruiqi Lin:** Data curation, Methodology. **Junchen Qian:** Data curation. **Yi Feng:** Methodology. **Shuang Chen:** Methodology. **Yuxin Wang:** Methodology. **Daoyou Chen:** Visualization. **Yanling Wu:** Project administration. **Tianlei Ying:** Project administration, Supervision, Writing – review & editing.

